# Th2 cytokine profile in appendicular lavage fluid is compatible with allergy as an etiology for acute appendicitis

**DOI:** 10.1101/569947

**Authors:** Nuno Carvalho, André Barros, Hélder O. Coelho, Catarina F. Moita, Ana Neves-Costa, Dora Pedroso, Filipe C. Borges, Luís F. Moita, Paulo M. Costa

## Abstract

Acute appendicitis is the most frequent surgical abdominal emergency, but its etiology remains poorly understood. Histological examination of the appendix, following its removal due to acute appendicitis, consistently shows features in common with bronchial asthma, suggesting an allergic reaction as a candidate etiologic factor. Here we propose the concept of appendicular lavage and used it to study the levels of the Th2 cytokines IL-4, IL-5 and IL-9 in patients with a clinical diagnosis of acute appendicitis. The study group included 20 patients with a histological diagnosis of phlegmonous appendicitis, 13 patients with gangrenous appendicitis, and a control group of 8 patients with clinical diagnosis of appendicitis but with normal histology. Cytokines levels were higher in acute appendicitis. The difference was more pronounced when comparing phlegmonous appendicitis with non-pathological appendix (p=0.01) for IL-4 (48,3 vs 21,3 pg/mL), IL-5 (29,2 vs 8,0 pg/mL) and IL-9 (34,1 vs 16,6 pg/mL). This Th2 cytokine profile is compatible with the hypothesis of allergy as an etiologic factor for acute appendicitis and may have important implications for the diagnosis, prevention and treatment of this condition.

## Introduction

Acute appendicitis (AA) is a frequent disease whose etiology cannot be explained by any single factor. Luminal obstruction is the trigger event that culminates in inflammation of the appendix. Fecaliths are found in one-third of specimens. In the other cases, obstruction is thought to be caused by hypertrophy of mural lymphoid follicles in response to diverse causes (1). The peak incidence of appendicitis coincides with the age when the immune response is most vigorous, and the lymphoid follicles are at their maximum development (2).

Aravandian has reported histological features in AA that are similar to bronchial asthma, a paradigm for an allergic reaction. Based on these findings, this author has proposed that AA is triggered by a type I hypersensitivity reaction and, therefore, could be caused by an allergic reaction (3). Cytokines from Th2 lymphocytes are responsible for the histological features of asthma (4). Th2 effector cells secrete mainly interleukin-4 (IL-4), IL-5, IL-9 and IL-13, which are known to be involved in allergic responses (5).

Broncho-Alveolar Lavage (BAL) is a useful tool for investigating inflammatory cell and mediator profiles, like cytokines, in various bronchopulmonary diseases (6). High levels of IL-4 and IL-5 are found in BAL in asthma (7). Similar to BAL, we have used the concept of appendicular lavage (AL), where saline is instilled and collected in appendicular lumen of appendectomy surgical specimens. The aim of this study was to test the hypothesis that AA might be the consequence of an allergic reaction by evaluating the levels of Th2 cytokines in AL fluid of patients submitted to appendectomy due to a clinical diagnosis of AA.

## Materials and Methods

### Study population

The study group, evaluated between April 2016 and June 2017, consisted of patients with the clinical diagnosis of AA, admitted to the emergency department of Hospital Garcia de Orta, when one of the authors (NC) was on call to perform the appendicular lavage. The only exclusion criterion was the absence of the author (NC). The histological diagnosis of AA discriminates acute phlegmonous appendicitis (APA) and acute gangrenous appendicitis (AGA). The control group consisted of patients admitted with the clinical diagnosis of AA, submitted to appendectomy, but with normal histology (Non-Pathological Appendix – NPA). No type of allergy test was performed. Laparoscopic appendectomy was performed in 33 patients and open surgery in 8 patients, including 3 conversions from laparoscopy. There were 9 localized and 3 generalized peritonitis.

### Ethics Considerations

This study is part of a research project approved by the Ethics Committee of Garcia de Orta Hospital (Reference 05/2015). Each enrolled subject gave written informed consent. The work has been carried out in accordance with The Code of Ethics of the World Medical Association (Declaration of Helsinki). The authors declare no conflict of interest.

### Appendicular lavage

After removal of the appendicular specimen, a gauge was inserted in the proximal luminal aspect of the appendix and 3 mL of saline 0,9 % were instilled and collected for AL. Saline was re-instilled and collected 3 times. Appendicular lavage process was standardized and performed exclusively by one of the authors (NC). The appendicular fluid samples were collected to a Sarstedt Monovette tube and centrifuged. 1 mL of the supernatant was extracted and stored at −20 °C. ELISA protocol was used for IL-4, IL-5, IL-9 and IL-6 determinations (Human IL-4, IL-5, IL-9 and IL-6 MAX, Biolegend, San Diego, CA 92121 USA) according to manufacturer’s protocol. The cytokine levels, IL-4, IL-5, IL-9 and IL-6 were expressed in pg/mL.

### Pathologic analysis

After AL procedures, the appendices were preserved in 10% formalin for histopathological examination. A minimum of 24 hours was allowed for adequate tissue fixation. Appendicular sections were sampled from the tip, base and intermediate length for fixation and paraffin processing. Two sections of 5-micron thickness were cut from each paraffin block and stained by hematoxylin & eosin (8). The criteria for AA was polymorphous nuclear neutrophils infiltration at the muscularis propria (9). APA was defined by the presence of neutrophils infiltrate in muscular propria and AGA was defined by the presence of necrosis of the wall of the appendix in a background of transmural inflammation (10). The presence of neutrophils in the mucosa was considered as a variant of normal with no clinical relevance, when no other inflammatory cells were detected in abnormal numbers. The specimens were classified as negative for appendicitis when no neutrophil infiltrate was show in muscular propria (NPA) (10). All the histopathological analyses were performed by one of the authors masked to the results of cytokine measurements (CH).

### Statistical analysis

Data are presented as descriptive statistics, mean and standard deviation. For continuous variables, considering the distribution of number of cases among the categories, a non-parametric approach was followed to assess statistical differences among the considered groups: Kruskal-Wallis tests were used, with *a posteriori* pairwise Wilcoxon tests; p-values were then corrected for multiple comparison using the Holm correction. For categorical variables, such as gender, a strategy based on Fisher’s exact test overall and pairwise, using the aforementioned correction, were used. Statistical Analysis was performed on R (https://cran.r-project.org), using the stats package for hypothesis testing and ggplot2 for the plots.

## Results

We analyzed 33 patients with a histological diagnosis of AA, 20 patients with APA, 13 patients with AGA and 8 patients, the control group, with normal histology. History of allergy was present in 7 patients (4 for antibiotics, 1 for metibasol®, and 2 with allergic rhinitis), no differences between groups were identified. None took medication. No differences in age, gender and BMI among groups were found (p=0.898, p= 0.054 and p=0.211, respectively) (Table 1). A significant difference was found for C-Reactive Protein levels and Length of stay (p=0.001, p= 0.002, respectively) (Table 1).

**Table 1.** Characterization of the population according to Histological Categories.

| **Variable** | **APA** | **AGA** | **NPA** |  |
| --- | --- | --- | --- | --- |
| **N (%)** | **20 (49)** | **13 (32)** | **8 (19)** | ***p*-value** |
| **Age (y)** | **36,2±16,9** | **20,0±15,1** | **34,2±12,9** | **0.898** |
| **Gender (M/F)** | **14/6** | **4/9** | **6/2** | **0.054** |
| **BMI** | **25,3±6,0** | **26,6±3,1** | **23,7±3,7** | **0.211** |
| **Allergy** | **3** | **2** | **2** | **NA** |
| **WBC** | **14,3±4,4** | **14,4±3,8** | **12,3±2,9** | **0,161** |
| **CRP** | **3,7±5,6** | **9,4±6,2** | **4,6±6,3** | **0,001** |
| **LOS** | **2,8±1,8** | **5,4±3,9** | **3,3±3,1** | **0,002** |

Results presented as number (valid percentage) or mean ± standard deviation. APA= acute phlegmonous appendicitis; AGA= acute gangrenous appendicitis; NPA= Non Pathological Appendix; M=Male; F=Female; BMI= Body Mass Index; NA= Not applicable; WBC= White Blood Count; CRP= C-Reactive-Protein; LOS= Length of stay.

The levels of cytokines in AL according to the histologic groups are presented in Table 2. In all the studied subjects, cytokines IL-4, IL-5, IL-9 were detected in AL fluid. For IL-4 there were significant differences among the histological groups (p=0.034). The difference between APA and NPA groups was significant (p=0.017). No significant differences for AGA group with the remaining groups were found (AGA vs APA p=0.421; AGA vs. NPA; p=0.421) (Figure 1). For IL-5, the differences were subtle (p=0.056). Differences among groups showed a tendency for different levels of IL-5 between APA and NPA (p=0.05) (Figure 2). As for IL-9, there was no clear evidence of differences for AGA group (AGA vs APA; p=0.587; AGA vs. NPA; p=0. 587). IL-9 was the cytokine whose levels revealed less differences among groups (p=0.083). However, while not statistically significant, there was a tendency indicating that differences exist between APA and NPA groups for this cytokine (p= 0.062) (Figure 3). No differences were founded between groups for IL-4 and IL-5 in the peripheral blood.

**Table 2.**
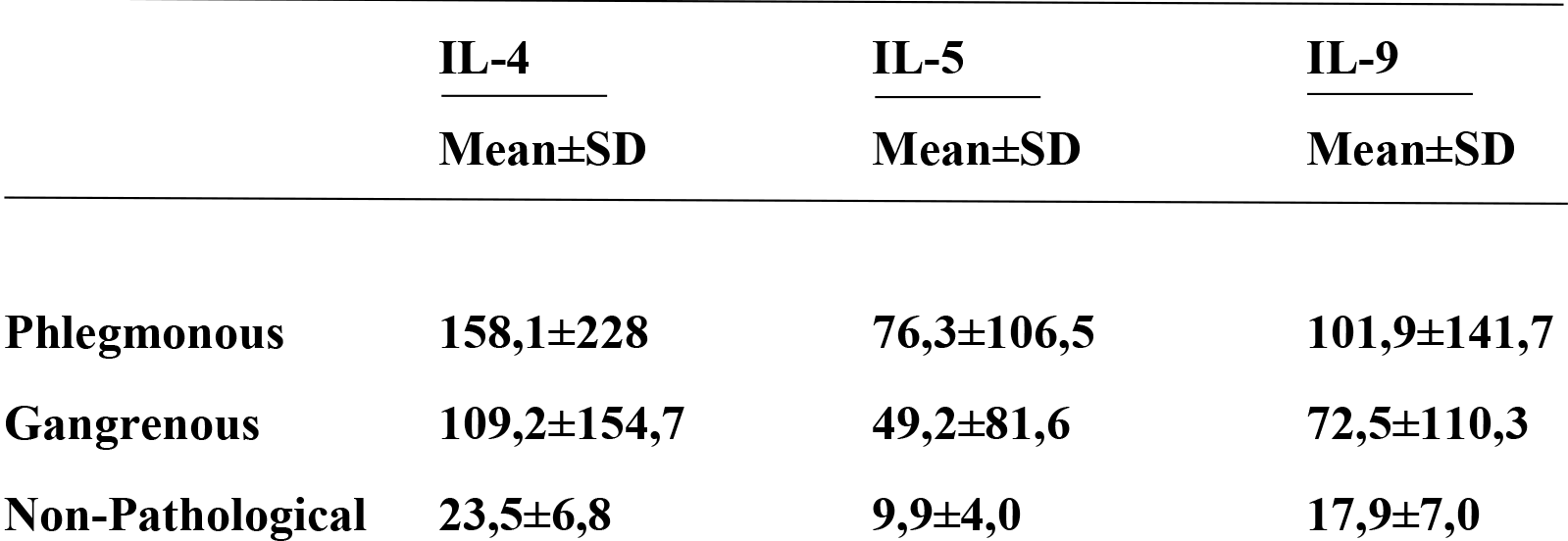
Cytokine levels (pg/mL) according to the histological categories.

**Figure 1.**
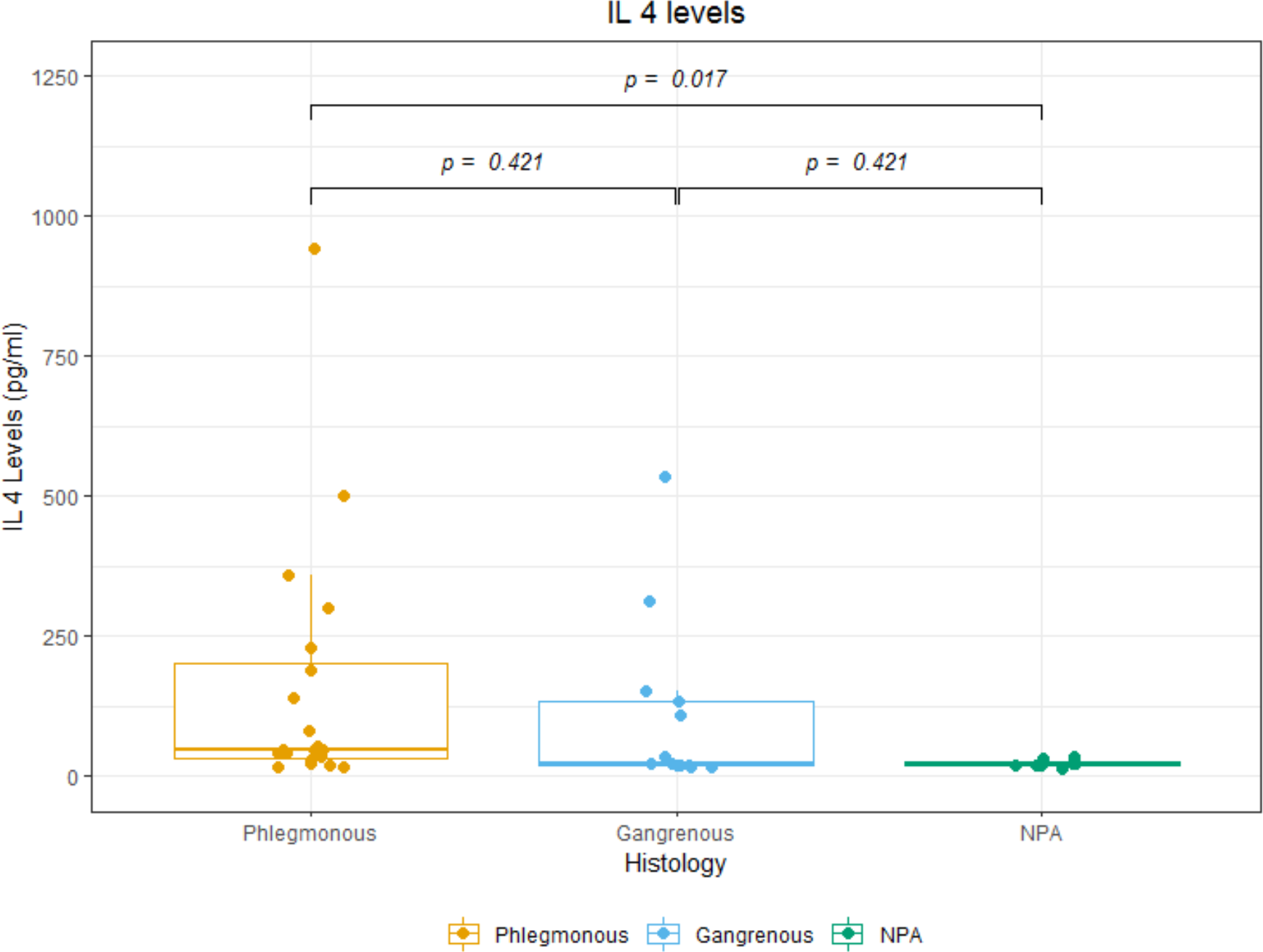
Box plots of IL-4 levels according to the different histological categories.

**Figure 2.**
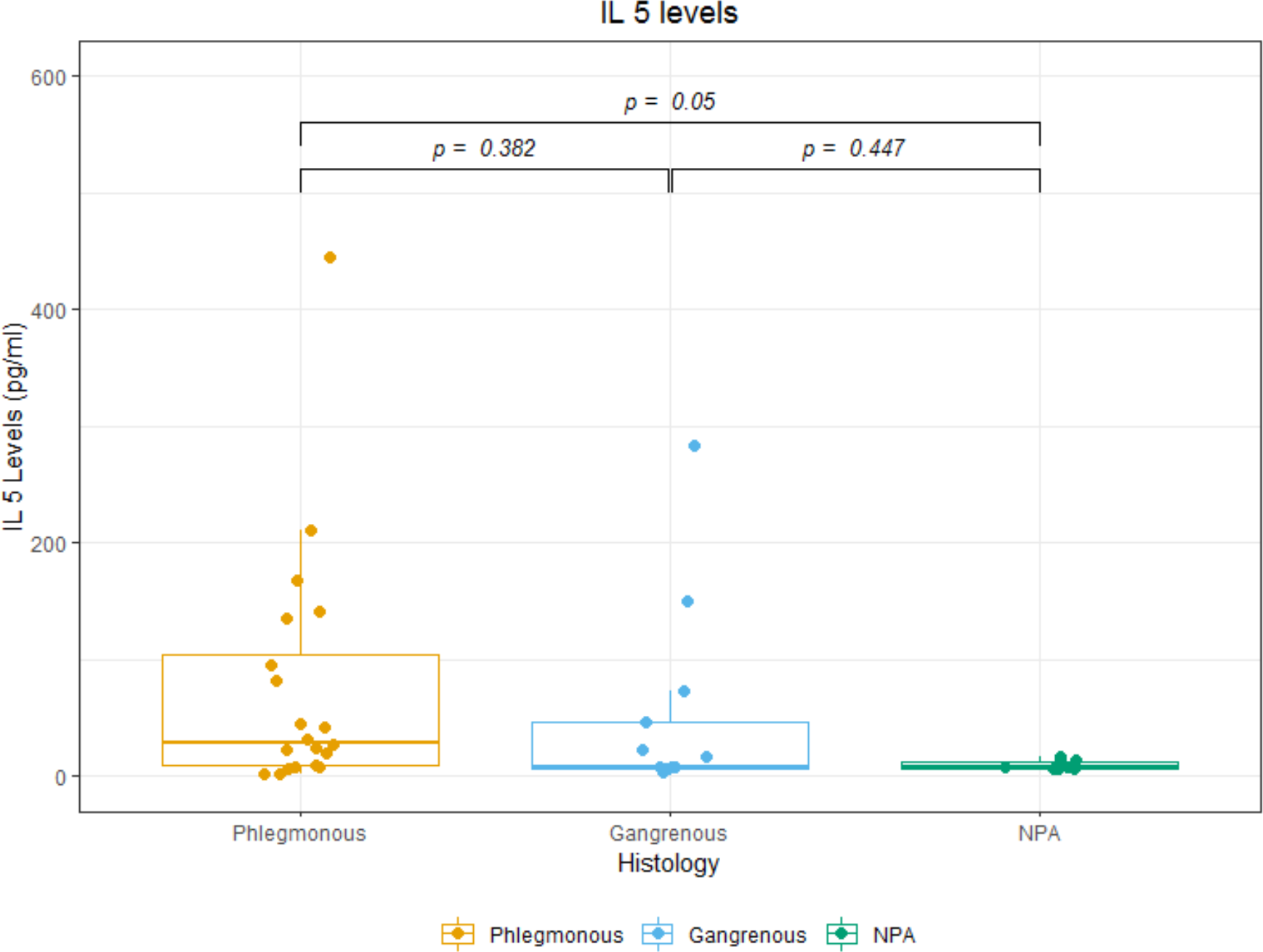
Box plots of IL-5 levels according to the different histological categories.

**Figure 3.**
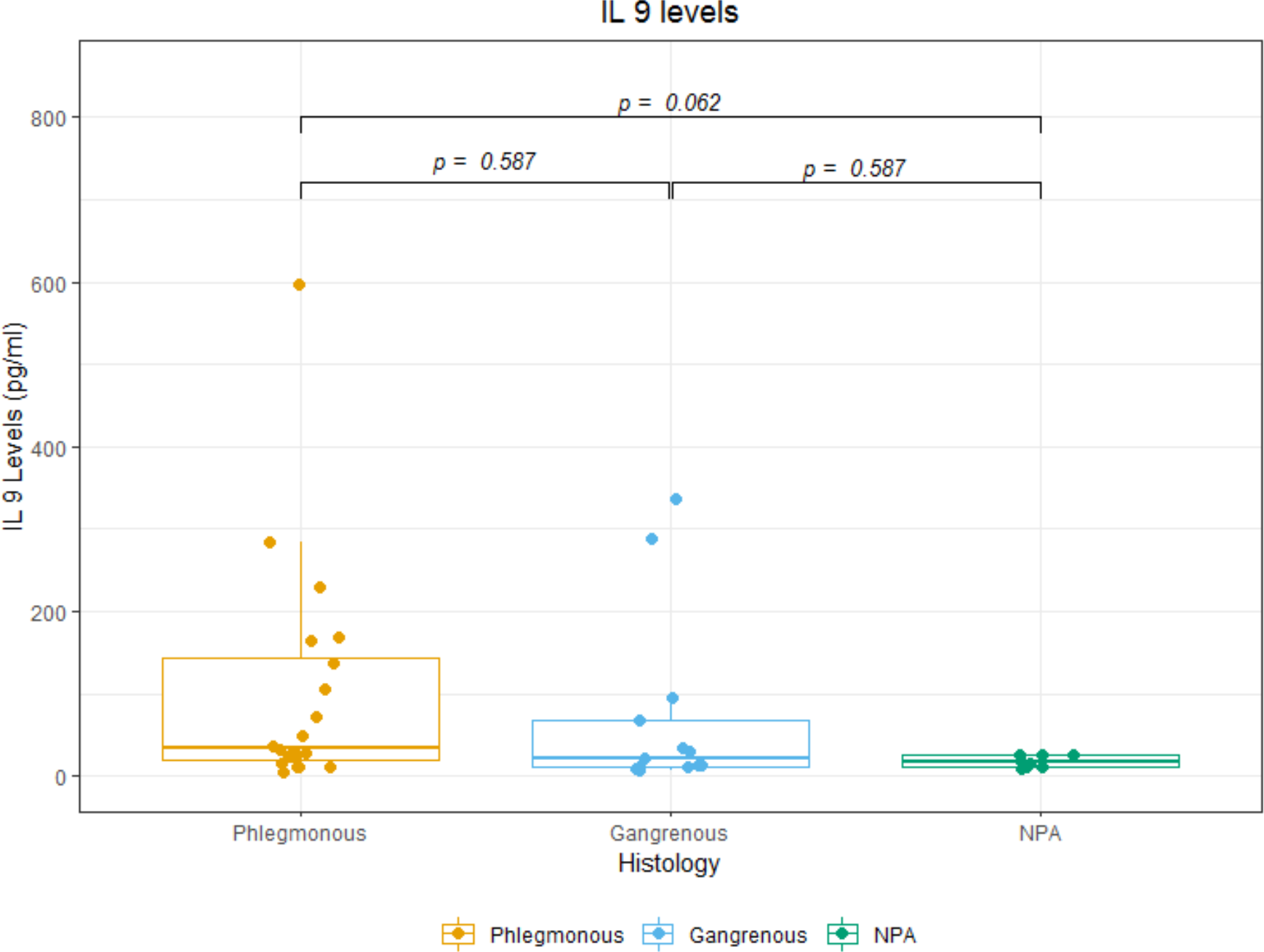
Box plots of IL-9 levels according to the different histological categories.

Th2 cytokines can be elevated in the course of an acute phase response together with other inflammatory mediators (11). To test whether Th2 cytokines were specifically associated with a histological diagnosis of AA, we have also measured the levels of IL-6 in the AL fluid, a prototypical general (non-allergic) acute phase cytokine (12). No differences were found among the groups (Supplementary Figure 1) in study (p=0.627).

To test an independent indicator of the contribution of an allergic reaction to the pathophysiology of AA, we have quantified IgE levels in the histological samples of the different groups (Figure 4). While differences among groups did not reach statistical significance (p=0.29) both the median (49 for APA, 42 for AGA and 26 for NPA) and mean (68.16 for APA, 48.85 for AGA and 23.50 for NPA) were substantially higher for APA and AGA when compared to NPA. These results, while not reaching statistical significance due to higher variation in the APA and AGA, strongly suggest that IgE levels are substantially higher in histologically documented cases of AA.

**Figure 4.**
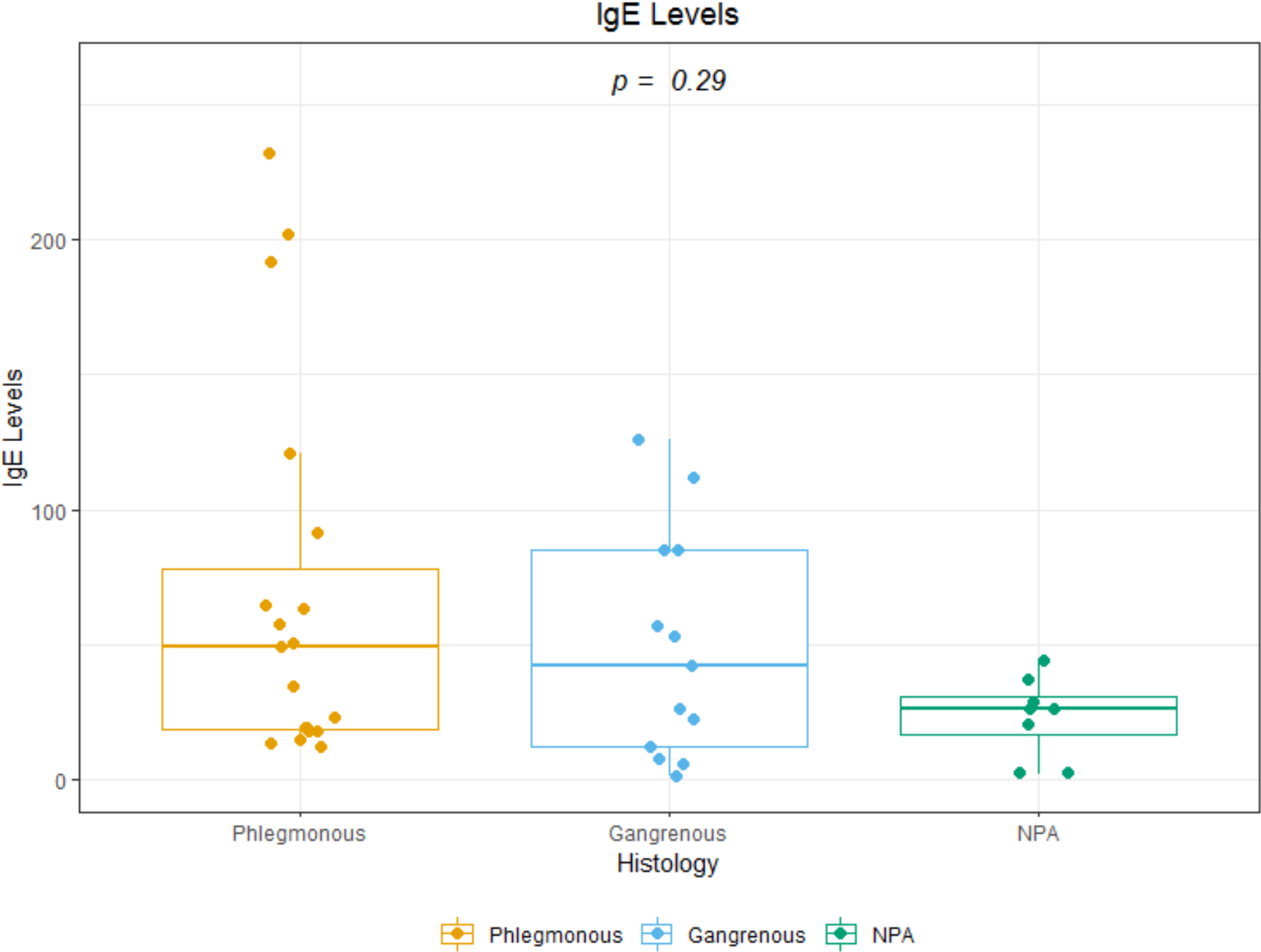
Box plots of IgE levels according to the different histological categories.

## Discussion

The most common theory for the etiology of AA is intraluminal obstruction, which is not supported by histological findings in surgical specimens. In most cases, no obstruction can be found (13). In one study, only in 0,4 % of 1969 appendectomy specimens was there evidence of luminal obstruction by vegetable fibers (14).

Based on histologic findings, specifically eosinophilic infiltration, mastocyte degranulation and muscular edema, an allergic reaction, was proposed as a possible etiology of AA (3). The concept is attractive: the appendix is a lymphoid organ, therefore, an immune response to a local antigen could be a factor in the pathogenesis of AA (15). The gastrointestinal system is one of the main entrances into the body for allergens during all life stages (9). Atopy may be a risk factor for appendicitis (9). By analogy with asthma, the contraction of the muscular wall of the appendix in response to antigenic stimulation, may result in luminal obstruction culminating in AA. This response may occur in any segment of the intestine, but the appendix is more vulnerable because of its small lumen size and limited capacity to accumulate fluids (3). The identification of the offending allergen(s) is not always straightforward (16).

Allergies are inflammatory diseases dependent on Th2 activation, mediated by IL-4, IL-5 and IL-9, in response to environmental allergens (7). IL-4 acts as a growth factor for Th2 cells and promotes the production of IgE. IL-5 induces the differentiation, activation and survival of eosinophils. IL-4 and IL-9 induce the growth of mast cells and basophils (7).

BAL has been performed for several years in lung diseases. We extend and adapt this approach to monitor the levels of cytokines in the appendix - appendicular lavage. The fluid collected from the appendicular lumen may reflect local inflammatory alterations. We used 3 washes with NaCl 0,9 % at the appendicular lumen to mobilize cytokines that are adherent to the mucosa. All steps of the process were standardized, from the surgical handling of the specimens to the processing of lavage and quantitative determinations. Differences in cytokines should be accepted as reflecting local inflammatory changes. Elevated levels of IL-4 and IL-5, are present in BAL of patients with allergic asthma (6). Higher levels are seen in symptomatic patients (17). The results of our study show a statistical difference in the levels of IL-4 between phlegmonous and non-pathologic appendicitis. IL-4 elevation reflects a putative allergic reaction in AA. In the case of IL-5, our data suggests differences between groups, mainly between phlegmonous and non-pathological appendix. For IL-9 there are also indications of differences between groups, mainly between phlegmonous and non-pathological appendix.

No significant differences for Th2 cytokines profile were founded between AGA and APA or NPA. In fact, some authors claim that simple inflammatory appendicitis and necrosis represent different diseases, or different patient response to disease, with distinct epidemiology, natural history, microbiology and different Th17 cytokine profile (18). Blood inflammatory response in AA showed a positive association of Th1 mediated immunity and gangrenous appendicitis (19). AGA etiology may be different from APA, without allergic component and so, Th2 cytokines are not elevated in AL. Another possible explanation for IL-4 and IL-5 similar values in AGA and NPA is that the cellular destruction in AGA is so marked that Th2 cells can no longer produce IL-4 and IL-5 and the values fall down to values found in NPA. This hypothesis is compatible with the possibility that AGA represents a later phase of the disease where tissue destruction is more pronounced and where the initial cytokines have been depleted.

In asthma, the paradigm of allergic disease, cytokine profile in BAL shows elevation of IL-4, IL-5 and IL-9 compared to control groups (6). Our results with AL are similar to those findings in BAL in the presence of allergy. In fact, our study showed elevations of Th2 cytokines in AL fluid of patients with appendicitis, compared with the control group (non-pathologic appendicitis).

### Strengths

Using a novel methodology, appendicular lavage, our study provides objective and original data demonstrating a correlation between cytokines and appendicular histological features that show that an allergic component is presented in AA.

### Limitation

Appendicular lavage should be reproduced by other researchers. The sample size is still limited. A larger dataset of patients, originating from multiple centers is strongly recommended because it is likely to validate and extend the conclusions of the current study.

### Conclusion

Differences in IL-4, IL-5 and IL-9 in AL fluid were found between the 3 study groups and especially significant between phlegmonous and negative appendicitis. Therefore, in AL fluid, we found a Th2 cytokine profile compatible with allergy. Further studies are required to assess the importance and robustness of our results regarding the allergic components in AA. The identification of an allergen will be a particularly difficult task, but our results open the possibility for novel management strategies in AA that might not always include a surgical procedure.

## Acknowledgements

We especially thank patients, surgical residents, surgeons, anesthesiologists, pathologists, nurses, medical and nursing students. Without their generous collaboration and hard work, this study would not have been possible. This research did not receive any specific grant from funding agencies in the public, commercial, or not-for-profit sectors.

## Author Contributions

N.D.C. designed the experiments, acquired data and wrote the manuscript. A.B.B., C.F.M. and A.N.C. did the analyses of the data. H.O.C. and F.C.B. acquired data including the pathology analyses of surgical samples. L.F.M and P.M.C. advised the project and reviewed the manuscript.

---

**Supplementary Figure 1.**
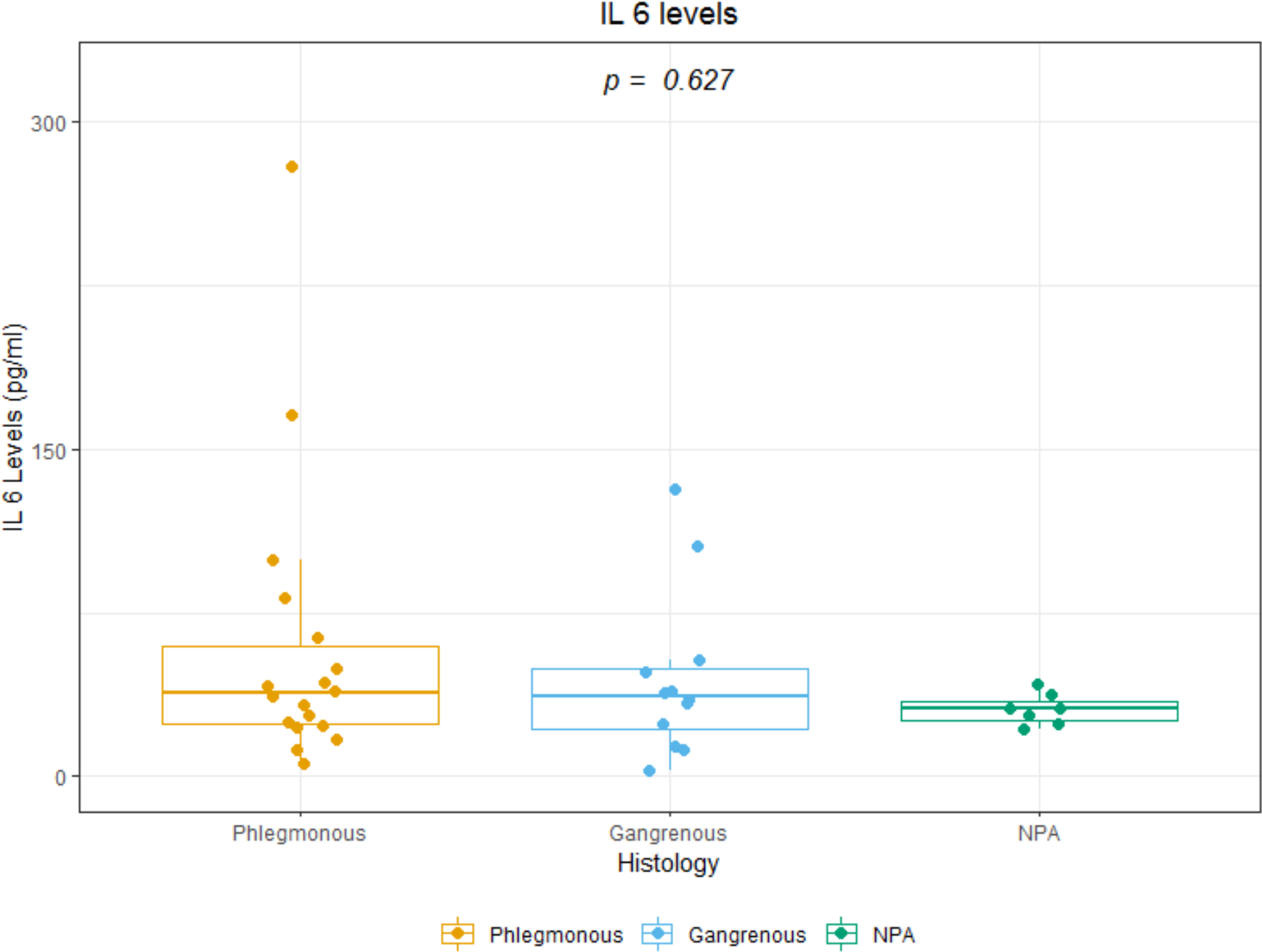
Box plots of IL-6 levels according to the different histological categories.

